# Efficient inference of recent and ancestral recombination within bacterial populations

**DOI:** 10.1101/059642

**Authors:** Rafal Mostowy, Nicholas J. Croucher, Cheryl P. Andam, Jukka Corander, William P. Hanage, Pekka Marttinen

## Abstract

Prokaryotic evolution is affected by horizontal transfer of genetic material through recombination. Inference of an evolutionary tree of bacteria thus relies on accurate identification of the population genetic structure and recombination-derived mosaicism. Rapidly growing databases represent a challenge for computational methods to detect recombinations in bacterial genomes. We introduce a novel algorithm called `fastGEAR` which identifies lineages in diverse microbial alignments, and recombinations between them and from external origins. The algorithm detects both recent recombinations (affecting a few isolates) and ancestral recombinations between detected lineages (affecting entire lineages), thus providing insight into recombinations affecting deep branches of the phylogenetic tree. In sim-ulations, `fastGEAR` had comparable power to detect recent recombinations and outstanding power to detect the ancestral ones, compared to state-of-the-art methods, often with a fraction of computational cost. We demonstrate the utility of the method by analysing a collection of 616 whole-genomes of a recombinogenic pathogen *Streptococcus pneumoniae*, for which the method provided a high-resolution view of recombination across the genome. We examined in detail the penicillin-binding genes across the *Streptococcus* genus, demonstrating previously undetected genetic exchanges between different species at these three loci. Hence, `fastGEAR` can be readily applied to investigate mosaicism in bacterial genes across multiple species. Finally, `fastGEAR` correctly identified many known recombination hotspots and pointed to potential new ones. Matlab code and Linux/Windows executables are available at https://users.ics.aalto.fi/~pemartti/fastGEAR/

## 1 Introduction

Microbial genomes are constantly subjected to a number of evolutionary processes, including mutation, gene gain and loss, genetic rearrangement and recombination, the latter here broadly defined as any form of horizontal transfer of DNA. The importance of recombination in prokaryotic evolution has been recognized for some time (Feil and Spratt, 2001; Didelot and Maiden, 2010; Hanage, 2016) and genomic studies have become an important source of data to measure its contribution (Polz *et al.*, 2013). Comparative studies of prokaryotic genomes have found that the vast majority of their genes have been laterally transferred at least once in the past (Dagan and Martin, 2007; Dagan *et al.*, 2008), with around 20% of genes being acquired recently (Popa *et al.*, 2011). Furthermore, when measured over shorter time scales, many bacterial species were found to recombine so frequently that the impact of recombination on their genetic diversification was shown to be greater than that of mutation alone (Vos, 2009).

The prevalence of recombination is suggestive of its importance for microbial evolution, with potential adaptive benefits. Genetic exchange between different strains has been argued to play an important role in shaping of bacterial communities (Polz *et al.*, 2013; Marttinen *et al.*, 2015; Shapiro, 2016) and the emergence of new bacterial species (Fraser *et al.*, 2007, 2009; Shapiro *et al.*, 2012). Bacterial recombination has also proved a powerful adaptive weapon against major forms of clinical interventions: antibiotics and vaccines (Hanage *et al.*, 2009; Croucher *et al.*, 2011; Perron *et al.*, 2012). As most evolutionary models (e.g., phylogenetic or phylogenomic analyses) assume no recombination, a good understanding of the impact of recombination on bacterial genomes is crucial for the correct interpretation of any genomic analysis.

Currently, popular methods used for detecting recombination in bacterial genomes include Clonal-Frame (Didelot and Falush, 2007) and ClonalFrameML (Didelot and Wilson, 2015), Gubbins (Croucher *et al.*, 2014b), and BratNextGen (Marttinen *et al.*, 2012). The three former approaches follow the line of methods based on phylogenetic trees (Husmeier, 2005; Minin *et al.*, 2005; Webb *et al.*, 2009), and look for clusters of polymorphisms on each branch of a phylogenetic tree. On the other hand, BratNextGen uses Hidden Markov Models (HMMs) to model the origin of changes in the alignment, where clonality (lack of recombinations) represents one origin and other origins represent foreign recombinations. All these methods specialise in identifying imports originating in external sources, and are therefore appropriately applied to a single bacterial lineage at a time. Thus, they rely on another method to identify the underlying population structure, which limits their ability to provide insight into species-wide or even inter-species patterns of exchange. With the recent development of high-throughput sequencing methods, which can process tens of thousands of bacterial whole-genomes, such analyses have become increasingly interesting and necessary.

Here we present an approach to fulfill such a demand, which identifies both the population structure of a sequence alignment and detects recombinations between the inferred lineages as well as from external origins. In particular, the method first identifies clusters of sequences that represent lineages in the alignment. Then it locates both recent recombinations, affecting some subset of strains in a lineage, and ancestral events between lineages that affect all strains in the lineage, as shown in Fig. 1. Our approach is similar to the popular STRUCTURE software with the linkage model (Falush *et al.*, 2003) but with the following crucial differences: (a) it is computationally scalable to thousands of bacterial genomes, (b) it provides insight into the mosaicism of entire bacterial populations by inferring also the ancestral recombination events. As our method can quickly infer the *ge*nomic *ar*rangement of large bacterial datasets, we called it `fastGEAR`.

**Figure 1:**
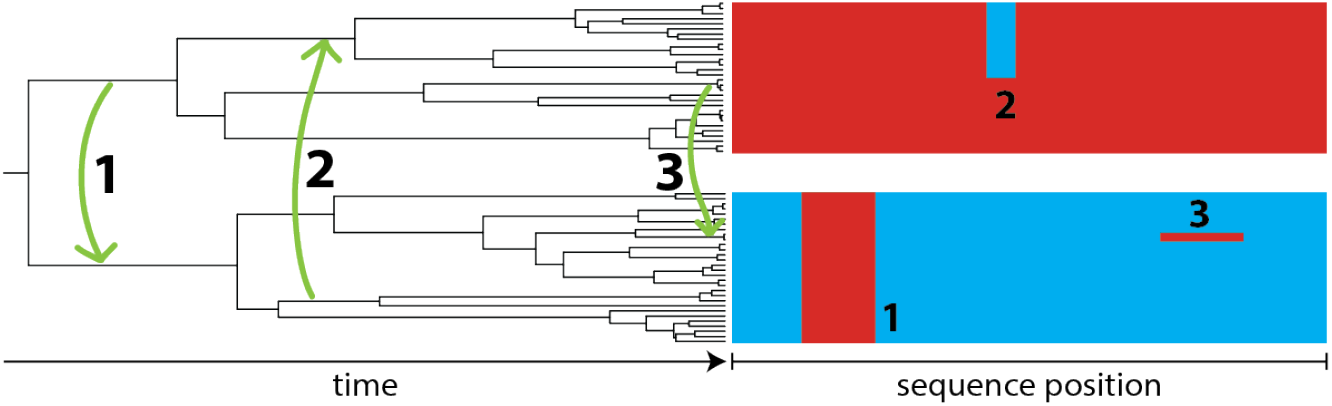
Simulations of bacterial recombinations.

The diagram shows the underlying simulation method, and here the case of *P* = 2 populations is considered: blue and red. Populations were simulated under a clonal model of evolution for a given set of parameters (see Methods). Three types of recombinations were then simulated using the clonal alignment. Ancestral recombinations (case 1) occurred before the most recent common ancestor of both populations, and thus were present in all isolates of the recipient lineage. Intermediate recombinations (case 2) occurred sometime between the time when populations emerged and present time (*t* = 0), and thus were typically present in multiple isolates. Recent recombinations (case 3) occurred in the last few generations, and thus were typically present in few isolates. To clarify, our method identifies events of type 1 as ancestral recombinations, whereas all other recombinations, affecting less than any whole lineage (cases 2 and 3), are inferred as multiple recent recombinations present in multiple individual strains.

We assess the accuracy of `fastGEAR` using extensive simulations, and compare it to three other state-of-the-art methods. We then use it to analyse a dataset of 616 whole-genomes of a recombinogenic pathogen *Streptococcus pneumoniae* sampled in Massachusetts, USA, in a paediatric carriage study (Croucher *et al.*, 2013). By applying `fastGEAR` to the assembled pneumococcal pan-genome, we obtain a high-resolution view of the underlying mosaicism of all coding sequences. We demonstrate that `fastGEAR` can be used on one hand for in-depth understanding of the extent of mosaicism in particular genes of interest (e.g., antibiotic resistance genes), and on the other hand to compare genome-wide the population structure and the extent of mosaicism in different bacterial genes. These analyses can be used in combination with commonly used pan genome assembly pipelines like Roary (Page *et al.*, 2015), thereby improving our understanding of the impact of recombination in bacterial genomes.

## 2 Overview of the approach

Here we give a general high-level description of the method, and the details are presented in Materials and Methods section. The algorithm takes as input an alignment of bacterial DNA sequences and performs the following four tasks:

1. Identify lineages in the alignment.
2. Identify recent recombinations in the lineages, where the ‘recent recombinations’ are defined as those that are present in a subset of strains in a lineage.
3. Identify ancestral recombinations between the lineages, where the ‘ancestral recombinations’ are defined as those that are present in all strains that belong to the lineage.
4. Test of significance of the putative recombinations.

The distinction between recent and ancestral recombinations is whether the recombination event happened before or after the most recent common ancestor of the lineage in which it was detected (see Fig. 1).

**(1) Identifying lineages** To identify lineages in a data set, we start by running a previously published clustering algorithm (Corander and Marttinen, 2006) included in the Bayesian Analysis of Population Structure (BAPS) software (Corander *et al.*, 2003). This produces C strain clusters which represent population structure among the strains. However, the clusters as such are not optimal for recombination analysis for two reasons. First, the algorithm may assign two otherwise identical sets of sequences into distinct clusters due to one of the sets experiencing a recombination event, because after the event the recombinant strains are more similar to each other than to strains in the other set, resulting in a suboptimal representation of the overall population structure across the sequence (see Fig. S1). Second, the algorithm may detect clusters that have diverged very recently, and the closeness of such clusters may result in added noise in recombination detection. Due to these reasons, given the clustering pattern, we infer lineages using a Hidden Markov Model (HMM) approach (Fig. 2A). In more detail, we compare allele frequencies for each cluster pair using a HMM, where the hidden states of the HMM represent equality of the frequencies at the polymorphic sites. All pairwise comparisons are summarised as a distance matrix, which tells the proportion of the length of the sequences where two clusters are considered different. We apply the standard complete linkage clustering with cutoff 0.5 to this distance matrix, resulting in a grouping of clusters into L groups, which are taken as lineages in the data. This means that two clusters will usually be considered as part of the same lineage if their sequences are considered similar for at least 50% of the sequence length, although this is not strictly enforced by the complete linkage algorithm.

**(2) Detecting recent recombinations** To identify recent recombinations, we analyse each lineage by applying a HMM approach for strains assigned to the lineage, this time with hidden states representing the origins of the different polymorphic sites in the strain (Fig. 2B). Possible origins are the other lineages detected in the data, as well as an unknown origin, not represented by any strain in the data. The positions which are assigned to a different origin than the identified lineage of the strain are considered recombinations. After analysing all strains in the lineage, the hyperparameters of the HMM are updated. Further iterations of detecting recombinations and updating hyperparameters are carried out until approximate convergence. The final reported recent recombinations are those sequence positions where the probability of the assigned lineage is less than some threshold, where we have used a conservative threshold value equal to 0.05. If a sequence position is considered recombinant, then the origin is set to be the lineage with the highest probability at this position. We note that also the full probability distributions are available from our implementation.

**Figure 2:**
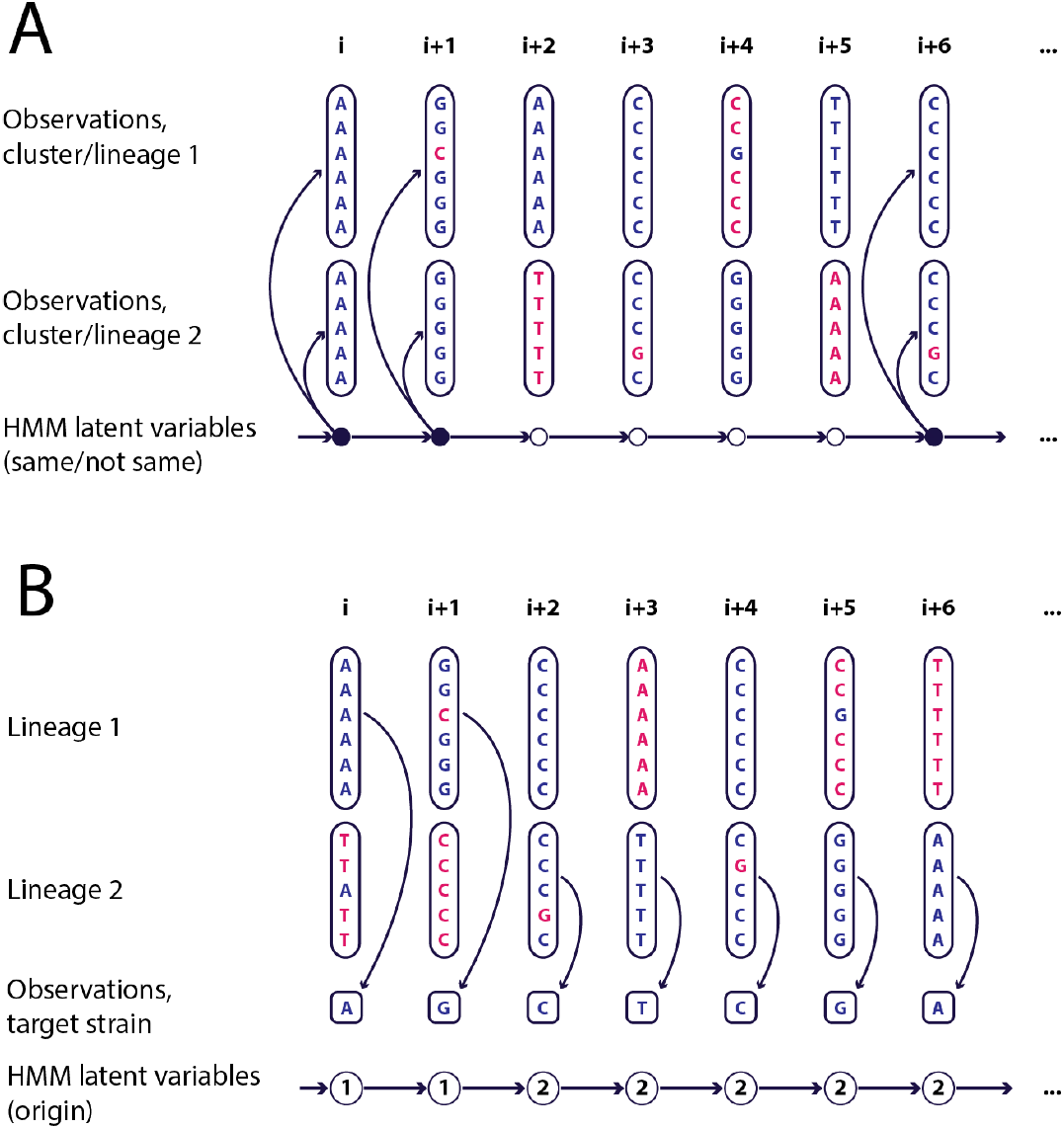
Hidden Markov models to detect recombination.

(A) Hidden Markov model used for identifying lineages and inferring ancestral recombinations. Each column represents a polymorphic site in the alignment and rows represent strains. The observed states of the chain are the allele frequencies within each cluster (in the case of identifying lineages) or lineage (in the case of identifying ancestral recombinations). The latent states of the chain represent identity of allele frequencies in the two lineages at the polymorphic sites. (B) Hidden Markov model used for identifying recent recombinations. The observed states are nucleotide values observed in the target strain and the latent states are possible origins of the nucleotides. The possible origins include all observed lineages plus an unknown origin.

**(3) Detecting ancestral recombinations** To identify ancestral recombinations, we analyse all lineage pairs using the same approach as in step (1), such that the latent variables for the different sequence positions have two possible states, either the lineages are the same or different, with the recent recombination fragments treated as missing data. Putative ancestral recombinations between lineages correspond to regions of the alignment where the inferred lineages are the same – hence a portion of the genome in isolates that are overall assigned to different lineages, may be considered to be part of the same lineage. However, it is important to note that the direction of a recombination can not be identified using this approach. To resolve this issue, we always mark the lineage with fewer strains as the recombination recipient in our results. The convention may be justified by the principle of maximum parsimony, as it results in fewer strains in the data set carrying a recombinant segment (but see Discussion).

**(4) Test of significance** The HMMs produce probabilities for sequence positions of having their origins in the different lineages, which can be used as a measure of statistical strength of the findings. However, in our experiments we encountered two kinds of false positive findings: first, recent recombinations in strains that were outliers in the data set; second, ancestral recombinations between lineages that were diverged to the verge of not being considered the same by the HMM, but not completely different either (see Discussion on the limitations of the HMMs). To prune these false positive findings, we monitor the locations of SNPs between the target strain and its ancestral lineage (for recent recombinations) or between the two lineages (for ancestral recombinations) within and between the claimed recombinant segments. We apply a simple binomial test to compute a Bayes factor (BF; see, e.g., Bernardo and Smith, 2001), that measures how strongly the changes in SNP density support a recombination, and we use a threshold BF=1 for recent recombinations and BF=10 for ancestral recombinations for additional pruning of recombinations proposed by the HMM analyses. These thresholds represent a compromise between false positive rate and power to detect recombinations. Recombinations with BFs less than the threshold are not reported at all, and the estimated BFs for the remaining recombinations are included in the output.

## 3 Results

### 3.1 Performance on simulated data

To give an example of `fastGEAR` performance, we performed coalescent simulations involving P = 3 lineages with a given effective population size for each lineage N_e_ and the most recent common ancestor (MRCA) time T. We then simulated recombination events between them in three different modes: recent, intermediate and ancestral. Finally, we compared the resulting, true population structure with the one inferred by `fastGEAR` in Fig. 3. We see that `fastGEAR` not only correctly identified the lineages, but also found almost all recombinations. The inference of recent recombinations was generally better than the inference of ancestral recombinations. Of particular importance is that `fastGEAR` does much better at predicting the direction of recombination events for recent recombinations. Such direction is difficult, if not impossible, to determine for older, ancestral recombinations. However, we see that the population genetic structure was correctly inferred in all three examples, even in the difficult case of multiple, overlapping recombinations occurring at different time scales.

**Figure 3:**
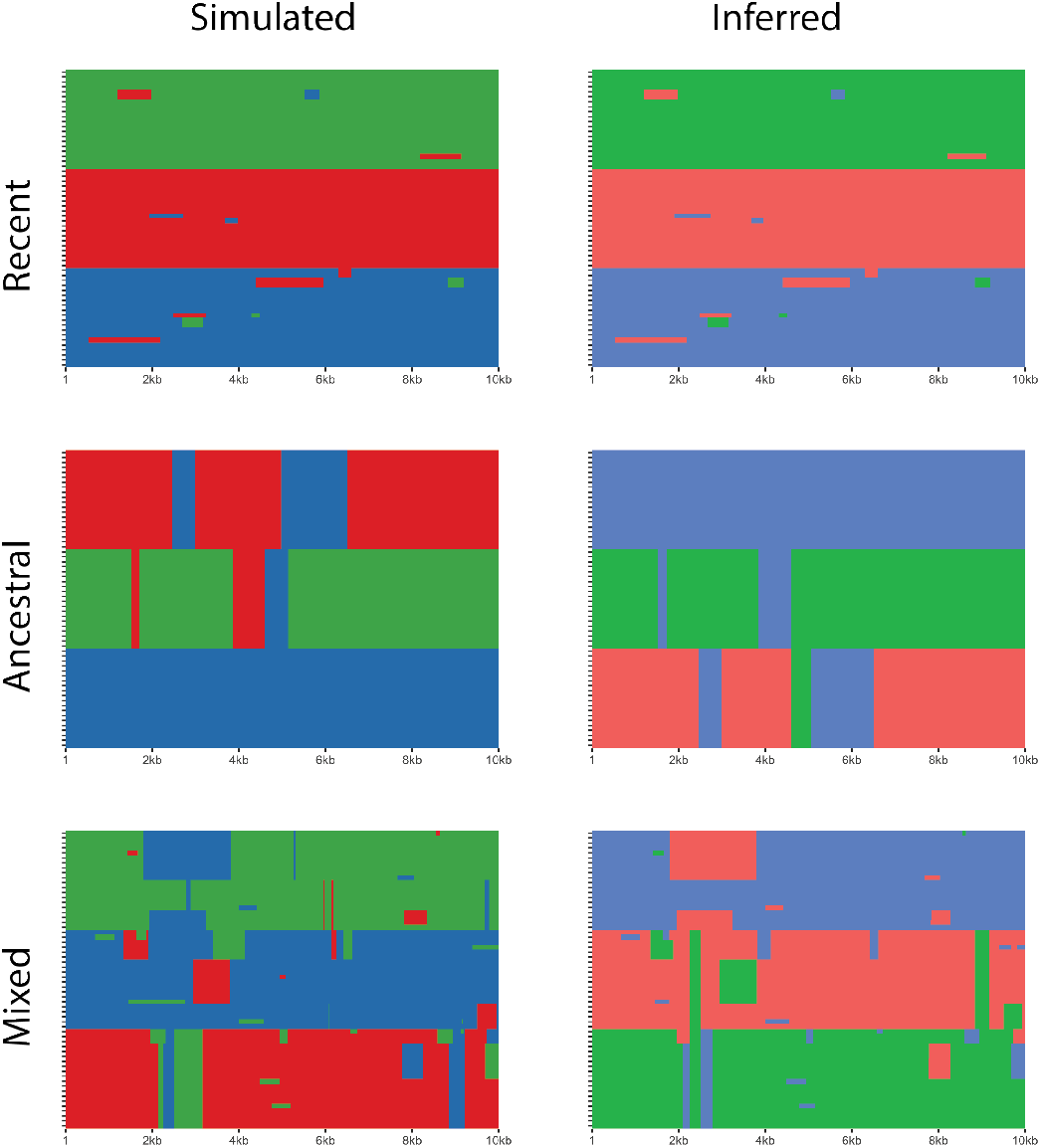
Visual assessment of the inferred population genetic structure.

The figure shows the population genetic structure of the simulated data. In each panel, the rows correspond to sequences, columns correspond to positions in the alignment and colours show different populations. The left column shows the simulated, true structure while the right column shows the population genetic structure inferred by `fastGEAR`. The order of the sequences in both columns is identical, and the colours are assigned randomly, thus populations are in the same order (1,2,3) but can be of different colour on the left and on the right. Three figure rows correspond to three different simulation scenarios: only recent recombinations (top), only ancestral recombinations (middle), and all three types of recombinations (bottom). The following parameters were used in the simulations: *P* = 3, *n* = 20, *N_e_* = 50, *T* = 2*e*4, μ = 2*e* — 6, *L* = 10kb (all rows); *Γ_r_* = 800 and *R_r_* = 5, *R_i_* = *R_a_* = 0 (top panel); *Γ_a_* = 800 and *R_a_* = 3, *R_r_* = *R_i_* = 0 (middle panel); *Γ_a_* = 500, *Γ_r_* = 500 and *R_a_* = 3, *R_i_* = 4 and *R_r_* = 6 (bottom panel).

To systematically assess the performance of `fastGEAR` we performed three different sets of *in silico* experiments. First, we examined how well `fastGEAR` detects recombinations for different population parameters. To this end, we varied the within-population distance (achieved by changing N_e_) and the between-population distance (achieved by changing T). The results are shown in Fig. S2. We see that `fastGEAR` generally detected recombinations well, particularly the recent ones as they share higher resemblance to the origin and are thus by definition easier to detect. The false-positive rate was low for all types of recombinations detected and did not vary with the between- or the within-lineage distance. By contrast, we observed that the proportion of detected recombinations was highly dependent on the between-lineage distance. This is because in the absence of clear population genetic structure, populations are relatively closely related and there are too few polymorphisms to signal the presence of a recombination. Furthermore, a higher within-lineage distance often affected the inference of ancestral recombinations as it generated the intra-lineage population genetic structure. Thus, as expected, performance of `fastGEAR` depends on the strength of the underlying population genetic structure.

In the second set of experiments, we compared `fastGEAR` to other recombination-detection methods: `STRUCTURE` (linkage model), Gubbins, and ClonalFrameML. To facilitate the comparison with the latter two phylogeny-based approaches, designed to detect recombinations from external origins in a lineage-by-lineage manner, and to explore the ability of `fastGEAR` to detect recombinations from unknown origins, we ran `fastGEAR` both using the entire alignment and with each lineage individually. Fig. 4 shows results in terms of false positive rate (the number of incorrect recombinations, left column), sensitivity (the proportion of true recombinations detected, middle), and the proportion of total length of true recombinations covered by detected recombinations (right).

**Figure. 4:**
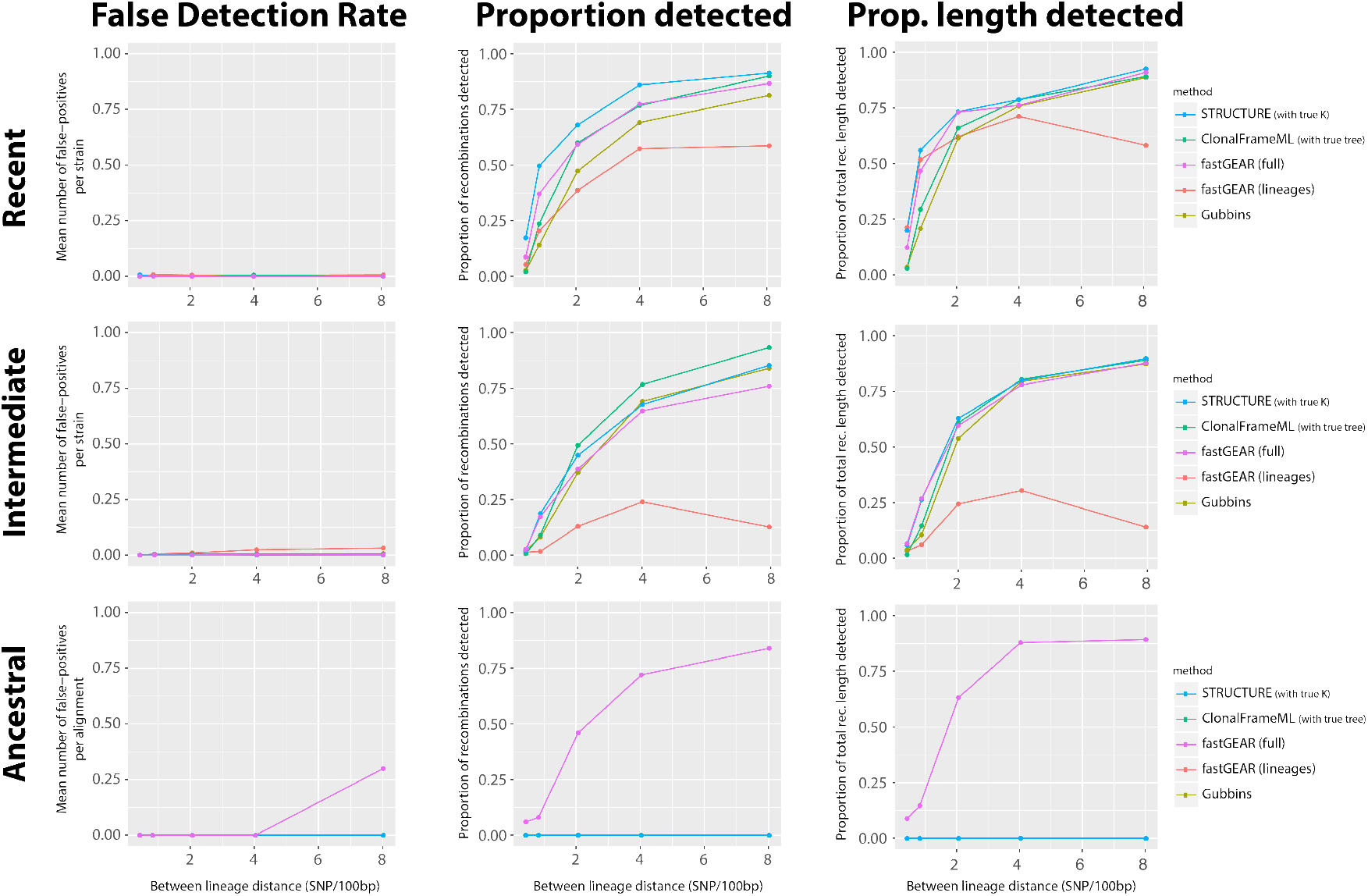
Comparison of fastGEAR and other recombination detection methods.

The figure shows the performance of `fastGEAR` compared to other methods: structure, Gubbins and ClonalFrameML. Top row shows results for recent, middle row for intermediate, and bottom row for ancestral simulated recombinations. Both recent and intermediate simulated recombinations were detected by `fastGEAR` in the same way as ‘recent’ recombinations. The left column shows the false detec-tion rate, namely, the mean number of false-positive recent recombinations per strain (top/middle) and ancestral recombinations per alignment (bottom). The middle column shows the proportion of true recombinations detected, and the right column shows the proportion of the total recombination length detected. Horizontal axis shows the between-population distance per 100bp (simulated by varying T between 1.0e3 and 2.0e4). Different lines show performance of different approaches. structure was run for 400,000 generations (200,000 burn-in), with true populations set as prior and with three independent chains to test for convergence. ClonalFrameML was conditioned on the true tree topology. Magenta line shows results of `fastGEAR` run on the full alignment, and red line for `fastGEAR` run lineage-by-lineage. Each point represents the average of ten simulations. The following parameters were used in the simulations: *P* = 3, *n* = 30, *μ* = 2.0*e*-6, *L* = 20kb, *Σ_a_* = 300, *Σ_a_* = 600 and *R_r_* = 5, *R_i_* = *R_a_* = 5.

We see that the false positive rate stayed very low with all methods. Furthermore, overall, `fastGEAR` had similar sensitivity to detect recent and intermediate recombinations to the other methods, and no method was systematically the best. Particularly interesting is the comparison between `STRUCTURE` and `fastGEAR`, because the two methods are generally quite similar, except that `STRUCTURE` explores the entire parameter space using MCMC whereas `fastGEAR` uses the most probable clustering and point estimates for hyperparameters. In general, `STRUCTURE` had a bit higher sensitivity, but at a much higher computational cost (between 1.5-2 orders of magnitude; see Fig. S3). Furthermore, `STRUCTURE` was here run conditioning on the true value of the number of populations K and the true lineage memberships of different strains as priors, while `fastGEAR` had no such knowledge.

When run in a lineage-by-lineage manner, `fastGEAR` had clearly lower sensitivity than with the full alignment, and the lengths of the external recombinations were often overestimated (see Fig. S4). This is because `fastGEAR`, similarly to `STRUCTURE`, gains its statistical power from having the actual origins of recombinations. Importantly, throughout simulations `fastGEAR` detected ancestral recombinations equally well to recent and intermediate recombinations. This is particularly encouraging as none of the other methods could detect ancestral recombinations. These results therefore show that while `fastGEAR` is able to detect external recombinations in a lineage-by-lineage manner, it is at its strongest when applied to investigate between-lineage or between-species bacterial data. An additional advantage of `fastGEAR` is that it can handle missing data in a straightforward manner, although, as expected, a lot of missing data may reduce the accuracy of recombination detection (see Fig. S5).

To facilitate the comparison of different methods, the previous benchmark simulations had predefined recombination probabilities that were not uniform in time, thus not allowing to investigate how the number of detected recombinations depends on some assumed fixed underlying recombination rate. To investigate this, we created a third set of experiments, where we simulated gene alignments from a single bacterial lineage with predefined internal and external recombination rates using ancestral recombination graph (ARG) simulations (Brown *et al.*, 2016). Fig. S6 and S7 show the results for different underlying mutation rates and gene sizes. Again, we see that the diversity levels affected recombination detection sensitivity, with a clear detection limit for extremely high recombination rates which reduced the linkage between SNPs to zero. On the other hand, we did not observe a significant effect of gene length on the detection rate: given a recombination rate, the number of detected recombinations per kb remained approximately constant, except for very short genes, where the length of a recombination may exceed the length of the gene, in which case the strains affected may be classified into a different lineage, rather than be identified as recombinant.

### 3.2 Analysis of *Streptococcus pneumoniae* data

We next demonstrate the utility of `fastGEAR` in three different data analyses using a whole-genome collection of 616 isolates of *S. pneumoniae.* First, we analysed alignments of three β-lactam genes responsible for penicillin resistance, augmenting the alignments with sequences from other species. This demonstrates the ability of `fastGEAR` to produce a detailed population structure of genome regions of interest and detect recombination between species. Second, we analysed the pneumococcal pangenome by running `fastGEAR` independently on the 2,113 codon alignments for COGs with at least 50 sequences from the previously-described pneumococcal population, to detect recombination hotspots in the pneumococcal genome. Third, we estimated a high-resolution genome-wide population structure of the pneumococcus population, by summarising the analyses of individual COGs in terms of the proportion of genome-wide shared ancestry between isolate pairs. The third analysis also demonstrates how the pneumococcal species-wide phylogeny, estimated from a core alignment, emerges as an average of highly variable population structures at individual loci, rather than representing clonal descent at any genome region.

#### 3.2.1 Inter-species recombination at penicillin-binding proteins

The population structure of *β*-lactam genes, which encode for penicillin binding proteins, has been previously analysed to understand the genetic basis of penicillin resistance and investigate the flow of antibiotic resistance determinants among streptococci (Croucher *et al.*, 2013; Jensen *et al.*, 2015). Here, we included the whole population of 616 *S. pneumoniae* strains, together with sequences from other species available in databases (list of accession numbers available in Table S1), resulting in the most comprehensive collection analysed for this purpose. We analysed each of the *β*-lactam genes, *pbp1a, pbp2b*, and *pbp2x*, using `fastGEAR`. We also investigated the sensitivity of the results on the presence of diverse strains by removing the outlier strains (i.e., those that belonged to lineages with five or fewer isolates in the original analysis).

Results with outliers removed are presented in Fig. 5, and they show a high-resolution view of the *pbp* genes across streptococci, with multiple recombination events between different species. Fig. S8 shows results before removing outliers, and we see that the inclusion of highly divergent outlier strains slightly lowers the sensitivity to detect recombinations, however the overall picture remains qualitatively identical. By comparison, Fig. S9 shows results separately for recent and ancestral recombinations, demonstrating that much of the structure remains hidden if ancestral recombinations are not considered.

**Figure 5:**
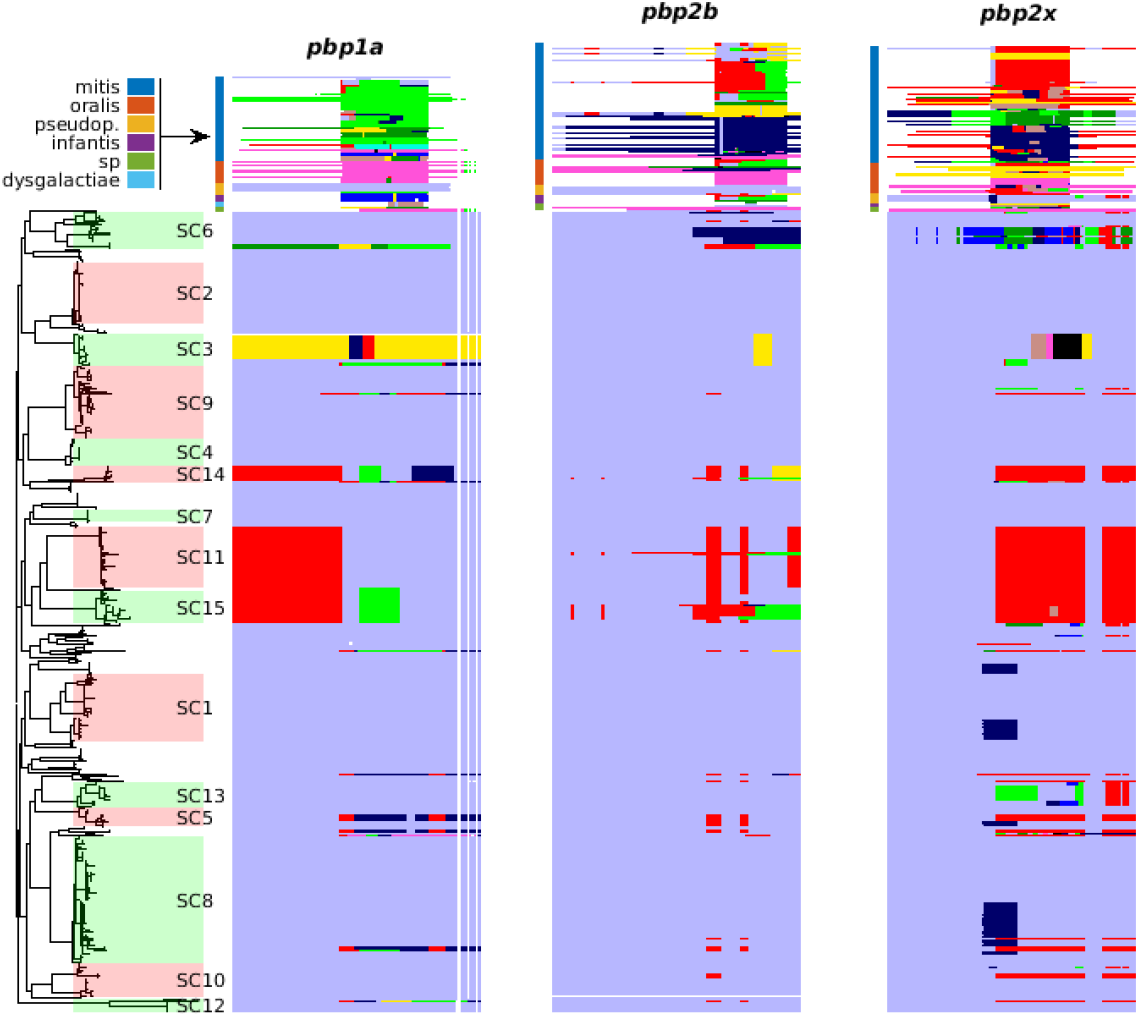
Results showing inter-species recombination at penicillin-binding proteins.

The figure shows `fastGEAR` results for combined data sets with the 616 S. *pneumoniae* strains and some number of additional sequences from other species (104 in *pbpla*, 129 in *pbp2b*, 127 in *pbp2x*). The phylogeny and the sequence clusters (SCs) on the left show the core-genome-based tree with 15 major monophyletic clusters for the *S.pneumoniae* strains. Strains from other species are shown on top of the *S. pneumoniae* strains. The species annotation is represented by colors on the left side of the additional strains, above the phylogeny. Note that the colors used to annotate species are independent of the colors in the `fastGEAR` output plots, where colors represent lineages detected by `fastGEAR`, except white which denotes gaps in the alignment.

Previously a software called BratNextGen (Marttinen *et al.*, 2012), developed by the same authors, has been used to investigate the population structure across diverse strains at the same loci (Croucher *et al.*, 2013; Jensen *et al.*, 2015). As mentioned previously, BratNextGen was designed for detecting imports from external origins in whole-genome alignments of closely related strains. For this reason, it does not automatically identify the origin of the imports, thereby requiring post-processing like manual curation in Jensen *et al.* (2015). To better understand how the difference between the two methods affects the inference of the underlying genetic mosaicism, we compared the results using both approaches using simulated data (Fig. S10) and the *pbp*’s (Fig. S11). The results show that the lineages detected by BratNextGen as well as the strains identified as recombinant are, broadly speaking, similar to those from `fastGEAR`. However, as expected, the BratNextGen output may be arbitrary if multiple diverse lineages are included into the analysis (Fig. S10). This can be best seen when comparing the results of the *pbp* analysis by `fastGEAR` and BratNextGen side-by-side, zooming in on the results across multiple species (Fig. 6; Fig. S12 shows results separately for recent and ancestral recombinations). One can see that `fastGEAR` identified mosaicism between species much better than BratNextGen. Overall, we conclude that `fastGEAR` is much more suited for the analysis of mosaicism of diverse bacterial genes than other, single-lineage methods like BratNextGen.

**Figure 6:**
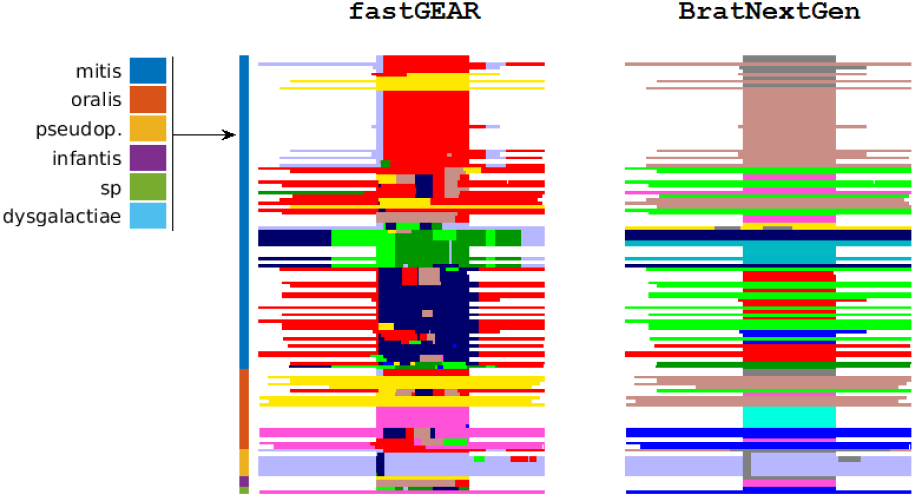
Comparison of `fastGEAR` and BratNextGen

The figure presents a detailed comparison of `fastGEAR` and BratNextGen. The results are shown for the *pbp2x* gene, zooming in to the sequences from multiple different species appearing on top in Fig. 5. We see that `fastGEAR` is able to detect mosaic structure between species.

We also investigated the relation between recombination and the levels of antibiotic resistance for *S. pneumoniae* isolates analyzed here (Fig. S13). The Spearman rank correlation between the detected recombinations and the minimum inhibitory concentration (MIC) was found significant in all genes (*pbp1a, ρ* = 0.66, p ≈ 0; *pbp2b, ρ* = 0.72, *p* ≈ 0; *pbp2x, ρ* = 0.75, *p* ≈ 0). Here the number of breakpoints due to both recent and ancestral recombinations was used as the variable tested against MIC, but all these correlations remained highly significant even if using either recent or ancestral recombination alone, or when representing the amount of recombination by the total length of recombinant sequence.

#### 3.2.2 Comparison of recombination levels across different proteins

We next compared the levels of recent and ancestral recombination between different proteins. Consistent with a constant rate of recombination over the history of this population, both measures were significantly correlated (R^2^ = 0.46, p ≈ 0) with the mean number of recent recombinations almost twice the number of ancestral recombinations (1.4 vs. 2.7; see Fig. S14). Based on the simulations, the overall diversity in a gene may affect the number of recombinations detected (see Fig. 4 and S6). To investigate the potential bias caused by this, we computed four common diversity measures and compared them with the estimated recombination counts (Fig. S15). No obvious relationship was seen, suggesting that results mainly represent true variation in recombination intensities.

Among the genes with the highest number of ancestral and recent recombinations shown in Tables 1 and 2, respectively, we found many loci previously identified as recombination hotspots (Supplementary Tables S2 and S3 show top hits for recombination intensities normalized by alignment length). These proteins can be classified into several groups. The first group consists of mobile genetic elements, which include integrative and conjugative elements, prophages, phage-related chromosomal islands and insertion sequences. This is not surprising as frequent between- and within-lineage recombination of mobile genetic elements has been reported previously (Croucher *et al.*, 2014a). The second group are proteins which are engaged in the interactions with the host. Pneumococcal surface protein C *(pspC*), which plays a central role in pathogenesis of the pneumococcus, was a top hit for both recent and ancestral recombinations (Kadioglu *et al.*, 2008). Another top hit was the first of the rhamnose genes *(rmlA)*, which often serves as a breakpoint in serotype switching events (see Supplementary text and Fig. S16 for a detailed discussion). We also found a high number of recent recombinations in the zinc metalloprotease *zmpA*, which cleaves human immunoglobulin A1 (Weiser *et al.*, 2003). The third group are genes involved in determining resistance to antibiotics, including sulphamethoxazole resistance *(folC*, see Fig. S17), as well as ß-lactams *(pbpla, pbp2b* and *pbp2x*) discussed above. With the exception of the mosaic *zmpA* sequences, these proteins were previously identified as recombination hotspots in globally disseminated lineages (Croucher *et al.*, 2011, 2014c).

**Table 1.** Summary of COGs with the highest number of ancestral recombination events (>20) identified by `fastGEAR`.

| SNPs | Sequences | Clusters | Lineages | Ancestral | Gene name |
| --- | --- | --- | --- | --- | --- |
| 788 | 288 | 17 | 10 | 58 | Pneumococcal surface protein C |
| 818 | 2149 | 24 | 17 | 51 | Insertion sequence IS1167 |
| 543 | 616 | 21 | 8 | 47 | Chromosome partition protein SMC |
| 618 | 2320 | 25 | 19 | 39 | Insertion sequence IS1381 |
| 297 | 325 | 14 | 11 | 34 | Bacteriophage DNA binding protein |
| 1177 | 378 | 10 | 7 | 32 | Tn5252 relaxase |
| 720 | 616 | 12 | 7 | 31 | Penicillin-binding protein 2x |
| 191 | 616 | 20 | 10 | 30 | Ribosomal protein L11 methyltransferase |
| 332 | 612 | 14 | 7 | 23 | Two component system histidine kinase |
| 258 | 336 | 11 | 7 | 23 | Capsule locus glucose-1-phosphate thymidyl transferase RmlA |
| 320 | 616 | 22 | 9 | 22 | Phenylalanyl-tRNA synthetase (PheS) |
| 310 | 229 | 11 | 7 | 22 | Protein found in phage-related chromosomal islands |

**Table 2.** Summary of COGs with the highest number of recent recombination events (>25) identified by `fastGEAR`.

| SNPs | Sequences | Clusters | Lineages | Recent | Gene name |
| --- | --- | --- | --- | --- | --- |
| 788 | 288 | 17 | 10 | 69 | Pneumococcal surface protein C |
| 543 | 616 | 21 | 8 | 69 | Chromosome partition protein SMC |
| 720 | 616 | 12 | 7 | 57 | Penicillin-binding protein 2x |
| 320 | 616 | 22 | 9 | 40 | Phenylalanyl-tRNA synthetase (PheS) |
| 536 | 616 | 16 | 5 | 37 | Valyl-tRNA synthetase (ValS) |
| 191 | 616 | 20 | 10 | 37 | Ribosomal protein L11 methyltransferase |
| 583 | 606 | 23 | 4 | 35 | Large cell wall surface anchored protein |
| 558 | 615 | 12 | 4 | 34 | Penicillin-binding protein 2b |
| 2021 | 248 | 10 | 4 | 34 | Zinc metalloprotease A |
| 490 | 616 | 11 | 7 | 31 | Folypolyglutamate synthase (FolC) |
| 263 | 614 | 17 | 7 | 31 | Transporter |
| 393 | 616 | 15 | 8 | 29 | Cell wall synthesis protein MurF |
| 618 | 2320 | 25 | 19 | 28 | Insertion sequence IS1381 |
| 326 | 597 | 15 | 5 | 28 | Choline binding protein C/J |
| 284 | 616 | 18 | 6 | 27 | Glucose-inhibited division protein GidA |
| 332 | 612 | 14 | 7 | 27 | Two component system histidine kinase |
| 741 | 97 | 7 | 5 | 27 | Phage protein |
| 574 | 614 | 9 | 5 | 26 | ATP-dependent protease ATP-binding protein ClpL |
| 242 | 613 | 19 | 5 | 26 | Mannosidase |
| 138 | 565 | 20 | 6 | 26 | Membrane protein |
| 421 | 64 | 9 | 6 | 26 | Part of large cell wall surface anchored protein PsrP |

We also found highly recombinogenic proteins which have not been previously identified as recombination hotspots. One example is the chromosome partitioning SMC protein, which functions in chromosomal segregation during cell division (Britton *et al.*, 1998) and is one of the top hits in both recent and ancestral recombinations. Other genes that were also inferred to undergo high levels of recombination are phenylalanyl- and valyl-tRNA synthetases, enzymes that attach the amino acids phenylalanine and valine to their cognate tRNA molecules during the translation process. Previous reports show that recombination and horizontal gene transfer frequently occur in aminoacyl-tRNA synthetases *(aaRS*) (Woese *et al.*, 2000). The horizontal acquisition of *aaRS* variants may be implicated in resistance to antibiotics (Woese *et al.*, 2000), and at least one atypical additional *aaRS* has been found on Pneumococcal Pathogenicity Island 1 (Croucher *et al.*, 2009). Although within-species recombination of *aaRS* has not been widely investigated, our results suggest that this process plays an important role in the evolution of pneumococci.

#### 3.2.3 High-resolution view of population structure

To investigate the population structure of the entire collection of isolates, we calculated the proportion of shared ancestry (PSA) matrix, which summarizes the `fastGEAR` results for all 2,113 COG alignments. Specifically, for each pair of isolates we analysed the population structure at each COG with putative lineages and the list of recent and ancestral recombinations detected in the COG. If the COG was present in both isolates, we computed the proportion of the length of the COG sequence in which the two isolates were assigned to the same lineage. If the COG was absent in either or both of the isolates, they were not compared at this COG; if multiple copies of the gene were found, then all possible comparisons between the two isolates were included, and correspondingly taken into account in the total length of sequence compared.

The resulting PSA matrix together with a previously published core-gene-based phylogeny and 15 monophyletic sequence clusters (SCs), which can be taken as lineages, is shown in Fig. 7. Overall, the PSA results are highly concordant with the tree and the SCs. First, strains within SCs share almost all of their ancestry, such that the average PSA within different SCs ranges from 85% up to 98%, which is visible as blocks of high PSA on the diagonal. Second, these blocks correspond well to the clades of the phylogeny. Third, the sequence cluster SC12, which has previously been identified as ‘atypical’, nonencapsulated pneumococci (Croucher *et al.*, 2014a) and appears distant from the rest of the population in the phylogeny, shares considerably less of its ancestry (approximately 60%) than other SCs share with each other. We also note that the polyphyletic SC16, which includes all strains in the phylogeny which are not part of SCs 1-15 (and is thus not shown), consists of multiple blocks of high PSA. These individual groups are similar to other SCs, with the difference that they are too small to be identified as separate SCs. Thus, we see that `fastGEAR` can produce a high-resolution view of the bacterial population genomic structure. Even though the PSA matrix and the phylogeny are in good accordance, our results highlight some details of the population structure not apparent in the phylogeny; for example, a pair of isolates between SC5 and SC8 in Fig. 7 that seem to share a large proportion of their ancestry with SC8.

A conspicuous feature of the PSA matrix is the lack of hierarchy between different SCs. Indeed, the different lineages (except for SC12) are approximately equidistant from each other, sharing from 71% to 81% of their ancestry, a pattern that can be explained by frequent recombination between the SCs (Fraser *et al.*, 2007; Marttinen *et al.*, 2015). To better demonstrate this, we computed the amount of private ancestry for each strain, defined as the proportion of the strain where the origin was not found in any other SC than the one to which the strain belonged (Fig. S18). The results show that all SCs have very little private ancestry; even the divergent SC12 has only about 15% of its ancestry private, i.e., not found in any other SC. These findings are consistent with the analysis of accessory genome content, which hypothesised that SC12 pneumococci may constitute a different streptococcal species altogether (Croucher *et al.*, 2014a).

To investigate the impact of recombination on the core genome further, we analysed the population structure of 96 housekeeping genes from an extended MLST set (Crisafulli *et al.*, 2013). The results for all the 96 genes are shown in Fig. S19, for a subset of 25 genes in Fig. 7, and for all core genes (i.e., present in at least 95% of the isolates) in Fig. S20. Two observations are particularly striking. First, we see that for the vast majority of genes the inferred number of lineages is much smaller than 15 (median: 3, 95% CI: 2-6). Second, the population structure is highly variable across the genome including at the 96 most essential genes, significantly deviating from a clonal model of diversification (Fig. S21 and S22). These findings lead to two important conclusions: (i) the pneumococcal-wide population structure, as represented by the SCs, emerges as the average of highly variable population structures of individual genes, and (ii) variable population structures of individual genes reflects their different evolutionary histories, and thus imply high rates of recombination at almost all bacterial genes, even the most conserved ones. These findings are consistent with an earlier comparison of phylogenies based on seven pneumococcal MLST genes (Feil *et al.*, 2001), and support the idea that recombination in some bacteria may eliminate the phylogenetic signal needed to establish relationship between different bacterial clones.

## 4 Discussion

In this article we introduced a novel tool called `fastGEAR` to analyse the population genetic structure in diverse bacterial isolates. Specifically, `fastGEAR` identifies major lineages and infers recombination events between them as well as those originating from outside the sampled population. Simultaneous inference of the population structure and between-population recombinations using Hidden Markov Models is analogous to an earlier approach called `STRUCTURE` (Falush *et al.*, 2003) but is novel in terms of both the ability to infer ancestral exchanges between those populations and being computationally scalable to thousands of sequences. In addition, our method is able to handle missing data in a straightforward manner, although for regions with lots of missing data the accuracy of the results may be decreased (see Fig. S5). Thus, our method is a notable addition to the currently available approaches for recombination detection in bacterial genomes, particularly so due to its ability to cope with increasingly large collections of whole-genome data and the ability to infer recombination in bacterial pan-genome data.

**Fig. 7.**
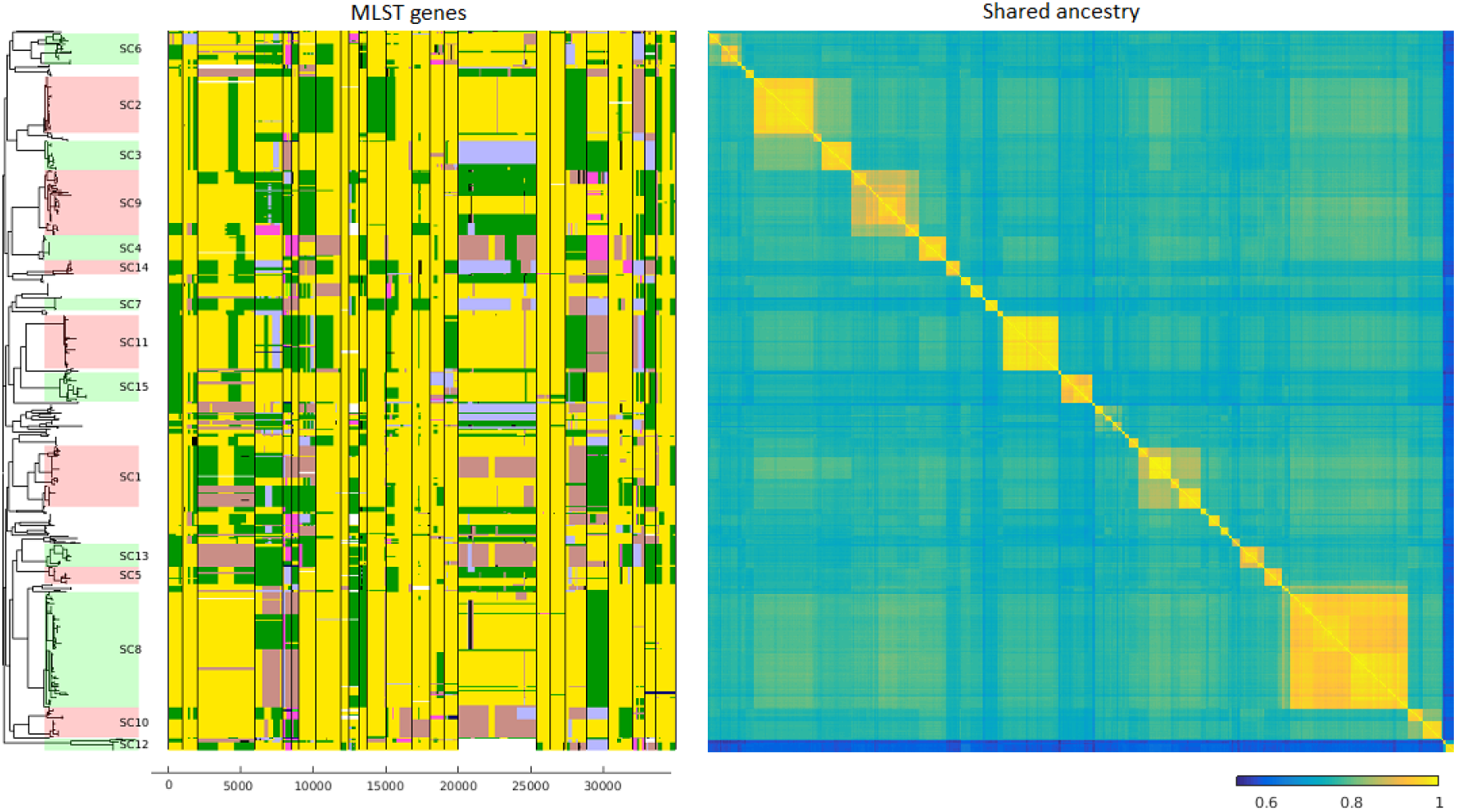
Population structure of the pneumococcal data.

The phylogeny and the sequence clusters (SCs) on the left show the core-genome-based tree with 15 major monophyletic clusters. Middle panel shows `fastGEAR` output for 25 out of 96 housekeeping genes, as discussed in the text; results for all 96 genes are qualitatively the same and shown in Fig. S19. The colors represent different lineages identified in the analysis (but are otherwise selected arbitrarily to be easily distinguishable). Recent and ancestral recombinations are colored with the color of the donor lineage. The results for the different genes were obtained by running `fastGEAR` independently, but the lineage colors at different genes were reordered to reduce the number of colors for any single strain across the genes (See Supplementary text for details). White colour denotes missing data. The PSA matrix on the right shows the genome-wide proportion of shared ancestry between the isolates in the data set, ranging from blue (distant) to yellow (closely related).

When detecting ancestral recombinations, our method is, broadly speaking, opposite to the clonal-frame-like approaches (Didelot and Falush, 2007; Croucher *et al.*, 2014b; Didelot and Wilson, 2015), where recombinations are divergent segments among highly similar sequences; here the ancestral recombinations are highly similar segments between diverse lineages. In this respect, our method resembles ChromoPainter/fineStructure (Lawson et *al.*, 2012), a method that investigates the similarity of haplotypes using other haplotypes as possible origins for the target haplotype, with the difference that `fastGEAR` detects recombination between groups of sequences. Our lineage-based approach has two benefits: first, if two or more strains share a recombination due to inheritance from a common ancestor, the recombination would not be seen at all if all sequences were considered as possible donors, because the recombination recipients are always closer to each other than to the donor. By considering lineages rather than strains as possible donors, it is possible to detect such a recombination. The second benefit is that in alignments with a very large number of highly similar or even identical sequences, identifying a specific donor would be very noisy, in particular because the actual donor is highly unlikely to be sampled in the collection, whereas identifying a lineage comprising a group of similar strains as a donor can be done robustly. The downside of the lineage-based approach is that recombinations between members of a single lineage are not detected. Our software implementation includes an option to manually define the lineages, which potentially could be used to detect recombinations between lineages defined at different resolutions, and our plan is to investigate this aspect in the future. Obviously, detecting recombinations between members of a single lineage is in practice limited by the lack of diversity between the strains, which reduces the power of any method to detect such events.

Even though `fastGEAR` detected ancestral recombinations exceptionally well in simulated data, a few points should be kept in mind when interpreting the results. First, the term ‘ancestral’ is relative and does not have to reflect the time of recombination; it merely reflects the fact that the recombination happened before the strains in the affected lineage diverged. Second, `fastGEAR` cannot reliably infer the direction of ancestral recombinations, as this would require additional assumptions about relationships between the ancestral sequences. We have resolved this by always marking the lineage with fewer strains as recombinant, assuming the lineage sizes are indicative of their ages, but the potential of sampling bias should be considered when interpreting results. This lack of directionality also means that while recombination breakpoints can be accurately inferred, determining conclusively which part of the sequence represents clonal inheritance (the clonal frame) is not possible. Therefore, interpreting the results in the phylogenetic context is recommended. We also note that a long ancestral recombination from a distant origin may appear in the results as a new lineage, rather than being reported as an ancestral recombination. Finally, we note that intermediate recombinations, present in multiple related isolates, are detected as multiple recent recombinations by the method. Therefore, to avoid reporting a single recombination event multiple times, the total number in the program output is obtained by considering spatially overlapping recombinations from the same origin as a single event (see Manual for further details).

Our statistical approach combines HMMs to identify putative recombinations with a post-processing step to compute the significances of the recombinations. These steps use information in the sequence data differently: the HMMs are based on allele frequencies at polymorphic sites, whereas the significances are computed using variations in SNP frequency along the sequence. The need for a separate post-processing step follows from the limitation of the HMMs that they can only tell whether two lineages are the same or different, but not how different they are. Consequently, very close or distant lineages are easily handled by the HMMs, but there always seems to be some intermediate distance for which HMMs may produce short segments of false positive recombinations, regardless of the exact way the HMM is formulated (for example, we experimented with various ways to handle the hyperparameters). The post-processing will produce a bias towards removing short, diverged segments as longer ancestral recombinations often reach higher significance. This is useful from a biological point of view because such short segments may also emerge as a combined result of mutation and positive selection. By assigning higher significance to longer fragments the chance of those fragments representing horizontal and not vertical evolution is increased. Nevertheless, a visual check of significance and a good understanding of the data analysed is always recommended.

Although our method does not assume a phylogeny, it nevertheless relies on some lineages between which recombinations are detected, and inferring the lineages is an important first step on which reliable downstream analysis can be based. Assuming consistent lineages over the length of one gene is more justified than over larger genomic regions, as demonstrated also by our results. For this reason we chose to analyse the *S. pneumoniae* data gene-by-gene, rather than concatenating multiple genes for joint analysis. The gene-by-gene analysis has additional benefits of being straightforward to parallelize and possible to apply to whole-genome core alignments. The latter is particularly appealing as it permits insight into the population structure and evolution of diverse microbial datasets, as well as analysis of standard bacterial pan-genome production pipelines. For these reasons, this is the way we currently recommend to use the method in practice. One downside of the gene-by-gene analysis is that there is no straightforward way for making inferences about long recombinations spanning multiple genes.

The usefulness of `fastGEAR` became evident when we applied it to a pneumococcal pan-genome from a whole-genome collection of 616 isolates from Massachusetts. By further inclusion of over 100 sequences from other bacterial species (including *S. mitis, S. oralis, S. pseudopneumoniae, S. sanguis* and *S. infantis)* to alignments of each of the three penicillin-binding protein loci *(pbp1a, pbp2x, pbp2b*), we gained a high-resolution view into the level of inter-species recombination within these important antibiotic-resistance genes. Furthermore, the analysis of all 2,113 genes produced a high-resolution view of the species-wide population structure. The population structure was consistent with previous studies of the fifteen major monophyletic groups but it also permitted insight into the ancestral composition of smaller clusters as well as to the relationships between the clusters. Finally, the analysis of recombinations within individual genes not only correctly identified many known major recombination hotspots in the pneumococcus but also pointed to potentially novel ones (SMC protein, *valS, aaRS*).

While developed and tested with bacterial genomes in mind, there is nothing in the method *per se* to exclude it from the analysis of other pathogens, including viruses. Nevertheless, `fastGEAR` assumes the isolates to be haploid, for which reason we expect `fastGEAR` to be particularly useful in questions related to microbial evolution. To conclude, `fastGEAR` offers a novel approach to simultaneously infer the population structure and recombinations (both recent and ancestral) between lineages of diverse microbial populations. We expect the method will bring novel insight into the evolution of recombinogenic microbial species, particularly so when recombination rates are high enough for the concept of species to be challenging to define.

## 5 Materials and Methods

### 5.1 Simulations

Details of simulations are given in Supplementary text. In brief, to generate *in silico* data we first created a phylogeny using a coalescent simulation framework Excoffier *et al.* (2013) assuming P = 3 demes which diverged T generations ago, each with a clonal population of effective size N_e_ and with mutation rate μ. A sample of n isolates was drawn from each population. An alignment of length L was created conditional on the phylogeny and recombinations were simulated by donating a homologous DNA fragment from a prespecified donor population to the target population, after which the fragment evolved according to the phylogeny of the target population. Recent recombinations were assumed to occur on average several generations before the present; intermediate recombinations were assumed to occur sometime between present and the youngest of all P most recent common ancestors for each population; ancestral recombinations were assumed to occur before the oldest of all P most recent common ancestors. The recombination size was modelled as a geometrically distributed variable with Г_г_ being the mean size of recent and intermediate recombinations and Γ_α_ the mean size of ancestral recombinations. We assumed on average R_r_ recombination events per population for recent recombinations (with targets chosen randomly), *Ri* recombination events per population for intermediate recombinations and *R_a_* recombination events for ancestral recombinations.

The accuracy of `fastGEAR` was assessed by quantifying the number of wrong recombinations (false-positives) and missed recombinations (false-negatives). To account for non-independence of recent and ancestral recombinations affecting multiple isolates, we clustered similar recombinations together with 95% identity threshold and counted each cluster as a single event. Inferred recombinations were then compared to true recombinations by comparing the isolates in which they occurred and position at which they occurred (assuming any overlap), which determined the number of false-positives and the proportion of all recombinations detected. Due to the difficulties in identifying direction of recombination, a detected recombination was considered a true-positive if the resulting population structure was correct, even if the recipient was not identified correctly.

### 5.2 Data from *Streptococcus pneumoniae*

We analysed a collection of 616 *Streptococcus pneumoniae* genome strains sampled in Massachusetts, for which whole-genome sequences were described in the original publication Croucher *et al.* (2013). The assembled data were scanned for putative protein-coding sequences, which were grouped according to their similarity, resulting in 5,994 clusters of orthologous genes (COGs). From these, we selected into our analysis those with at least 50 sequences, and we only included proteins that were within 75% and 125% of the median length of the COG. After this filtering, we kept COGs with at least five distinct protein sequences included in the data set, resulting in a total of 2,113 COGs included into our analysis. Unique sequences were aligned with Muscle Edgar (2004). All DNA sequences associated with each protein sequence were then back-translated into a full codon alignment. A core alignment was constructed using COGs present once in each genome assembly. This alignment was previously used to produce a maximum likelihood phylogeny of the data, and analysed by BAPS to produce 16 sequence clusters (SCs), of which 15 were monophyletic Croucher *et al.* (2013).

### 5.3 Details of the algorithm

#### 5.3.1 Identifying lineages

Let *X* = [*x_ij_*] denote the alignment of polymorphic positions *j* = 1,…, *J* in strains *i* = 1,…, *N*. We use the BAPS algorithm (Corander and Marttinen, 2006) to estimate a partition 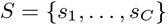 of strains into C separate clusters *s_c_*. To identify lineages in the data, a HMM is used to compare all cluster pairs. To define a HMM, a probability distribution for the first latent variable, *z_1_*, a transition matrix, and emission probabilities must be specified (see, e.g., Bishop, 2006, Ch.13). We use a uniform prior

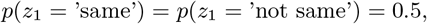

and we define a transition matrix *T_j_* for a transition from the (*j* — 1)^*st*^ to the *j^th^* polymorphic site as

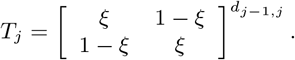

Here *ξ* is a hyperparameter that specifies the probability that if the clusters are considered the same in some sequence position, they are the same in the next position as well. The distance *d_j-1,j_* between the polymorphic positions is taken into a account by raising the matrix to the power *d_j-1,j_*. We use a fixed value for the hyperparameter

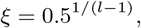

which corresponds to the assumption that the two clusters are the same for the entire length *l* of the sequence with probability 0.5. We note that when comparing clusters (and later lineages) using the HMM, the emission probabilities, which are based on all observations in the clusters, are highly informative, and therefore the conclusions are insensitive to the exact value of the parameter ξ; replacing 0.5 by 0.05 or 0.005 produced almost identical results in our experiments. Let 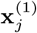 and 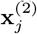 denote observations at locus *j* in the two clusters that are compared, and let 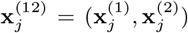. We specify the emission probabilities as

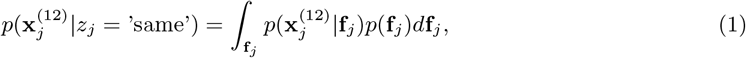

where 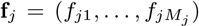 are the frequencies and *M_j_* the number of the different alleles observed at locus *j*. Furthermore,

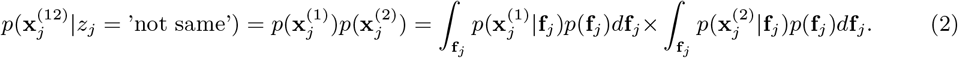

The likelihood in (1) and (2) is defined as

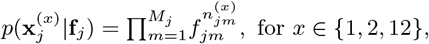

where we have denoted the count of allele *m* at locus *j* in group *x* by 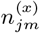. The prior on the frequencies *f_j_* is defined as

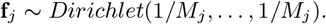

With this prior it is possible to integrate out the frequencies analytically, which gives

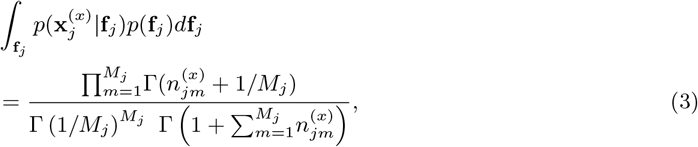

which can be used to compute (1) and (2).

Finally, the HMM analysis is used to calculate a distance matrix between the clusters *s_c_* detected in BAPS analysis, with the distances representing the proportion of sequences considered ‘not same’ by the HMM. This distance matrix is used as an input to the standard complete linkage clustering, and lineages are defined by using a cutoff 0.5, resulting in another partition 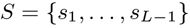 of the strains into *L* — 1 lineages. We add to this an empty lineage 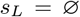 that represents an unknown origin of recombinations, not represented by any strains in the data.

#### 5.3.2 Detecting recent recombinations

To detect recent recombinations in a lineage, we analyse all strains in the lineage with a HMM, and repeat this for all lineages. We call the lineage that is currently analysed the *ancestral lineage* for strains that belong to the lineage, while other lineages are called *foreign* for the strains. The strains in the lineage are analysed consecutively, after which the hyperparameters of the HMM are updated, after which the strains are again analysed, and this is continued until approximate convergence. We note that we use a separate probability model for each lineage as opposed to a more formal approach with a joint model over all data. This simplification enables computation that is scalable up to thousands of strains in dozens of lineages.

The HMM is defined with transition probabilities that represent switches in the origin of the target strain between the ancestral lineage and foreign lineages. We define these in a way that resembles the transition probabilities used in the BratNextGen (Marttinen *et al.*, 2012), with the difference that in BratNextGen the transitions happen between a single non-recombinant state that represents the clonal ancestry of all strains in the data and other states that represent possible recombination origins outside of the data. Let *l_anc_* denote the lineage that is currently analysed. Then the transition matrix at a polymorphic position *j* is specified as

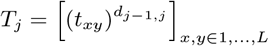

where

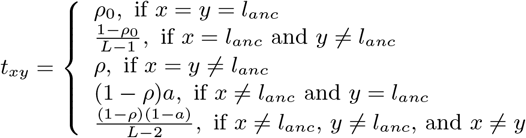

The different parameters are interpreted as follows: *ρ_0_* is the probability of staying in the ancestral lineage; 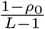 is the probability of moving away from the ancestral lineage into a foreign lineage, ρ is the probability of staying in a foreign lineage; (1 — *ρ)a* is the probability of moving from a foreign lineage back to the ancestral lineage; 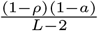 is the probability of moving from a foreign lineage to a different foreign lineage.

Emission probabilities are defined by assuming that 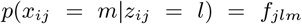, where 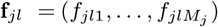 are the frequencies of the different alleles at locus j in lineage l. With the prior 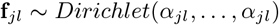, we can integrate out *f_jl_*, which yields the emission probabilities

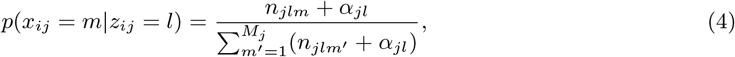

where *n_jlm_* is the count of allele *m* at locus *j* in lineage *l*. When calculating *n_jlm_*, we remove the target strain from its ancestral lineage, as well as recombinant regions of other strains that belong to the lineage. Also, for the external origin, corresponding to *l* = *L, n_jlm_* = 0 always. An important consideration is how to specify the hyperpameters *α_jl_.* One possibility would be to set *α_jl_* = *c_j_*, with some *c_j_* that would be constant across the lineages. This has the downside that if sequences in multiple lineages are exactly the same, then the origin corresponding to the largest lineage would have the highest probability according to formula (4). However, we want the model to assign the highest probability to the ancestral lineage unless there’s evidence against it, which we achieve by assuming overdispersion in the multinomial distributions of the allele counts (see, e.g., Poortema, 1999) in those foreign lineages that are larger than the ancestral lineage. In practice, we specify

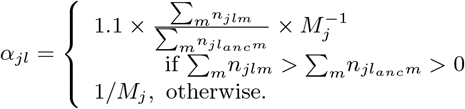

This ensures that the ancestral lineage will always have a higher value of (4) than a foreign lineage, even if all sequences in both lineages are identical to the value in the target sequence. Finally, a unifrom prior is again used for the first latent variable, completing the definition of the HMM for recent recombinations.

#### 5.3.3 Detecting ancestral recombinations

Ancestral recombinations between lineages are estimated by first removing the recent recombinations in the lineages detected in step 2, and then analysing each lineage pair using a HMM similar to the one that was used for comparing different pairs of clusters when estimating lineages in step 1. This tells us sequence regions in which two lineages are the same (with probability >0.5) or different. We note that the direction of recombination is not identifiable by this approach, it only tells whether two lineages are ‘the same’ or ‘different’ in some region. Therefore, after we have computed the similarity patterns for all lineage pairs, we follow the convention stated in the main text, and assign the regions of the smaller lineage to have their origin in the largest lineage that is similar to the smaller lineage in the region.

#### 5.3.4 Test of significance based on SNP density

To compute an additional significance check for putative recent recombinations, we identify locations of SNPs between the target strain and the summary sequence (modal sequence after removing recent recombinations) of the ancestral lineage of the strain. The SNP density between the non-recombinant regions is compared with SNP densities within the putative recombinations. In detail, let 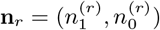 be the numbers of SNPs and non-SNPs within the *r^th^* recombination and 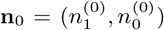 the corresponding numbers in the non-recombinant regions. We compute a Bayes factor (BF)

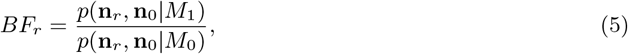

where *M_1_* assumes that *n_r_* and no come from different distributions

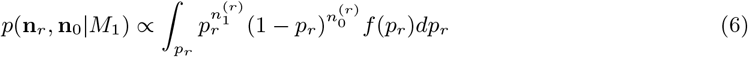

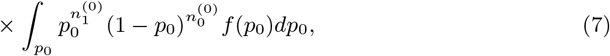

where *p_r_* and *p_0_* denote SNP frequencies within the recombination and non-recombinant regions, respectively. *M*_0_ assumes that the observations come from the same distribution with a single frequency parameter *p*

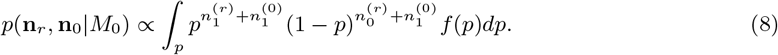

We note that the proportionality constants in (7) and (8) are equal and, thus, cancel when computing (5). With a non-informative uniform prior 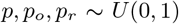, the integrals in (7) and (8) can be computed and equal

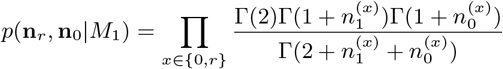

and

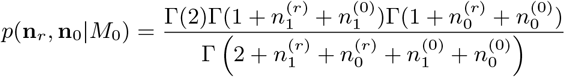

In detail, the algorithm continues as long as there are recombinations with *BF* less than 1, and always removes the recombination with the lowest BF, combines observations from the removed recombination with the non-recombinant observations, and recomputes the Bayes factors for all remaining recombinations.

We compute analogous Bayes factors for the ancestral recombinations. In detail, we process the lineages one by one, such that we consider the segmentation of the lineage into different origins. For each segment, we compute the Bayes factor using a similar beta-binomial model based on changes in SNP density between segments. SNP density is here computed between the origin of a recombination and non-recombinant regions (for recombinant segments), or between the ancestral lineage and origins of its neighboring segments (for non-recombinant segments). If the BF for SNP density changing is less than 10 for any segment, then the segment with the smallest BF will be combined with its neighbor, such that always the shorter segment is merged with the larger one. The thresholds for recent and ancestral recombination BFs, 1 and 10, correspondingly, were selected based on preliminary experiments, and represent a compromise between false positive rate and power to detect recombinations.

## 6 Supplementary Material

Supplementary material consists of a single PDF containing Supplementary text (simulation setup, reordering of lineage colors, detailed results for clinically relevant genes), Fig. S1-Fig. S22, and Tables S2-S3. Supplementary Table S1 is uploaded separately as an XLSX file.

## Acknowledgments

This work was funded by the Academy of Finland (grants no. 286607 and 294015 to PM) and Junior Research Fellowship from Imperial College London (RM). The calculations presented above were performed using computer resources within the Aalto University School of Science “Science-IT” project and computing systems at the Wellcome Trust Sanger Institute. Authors would like to thank Xavier Didelot for helpful comments on the manuscript, David Aanensen and Yonatan Grad for insightful discussions, as well as Stephen Bentley and Julian Parkhill for kindly providing access to the Sanger computational resources.

